# The Mechanisms Underlying Colour Afterimages

**DOI:** 10.1101/2022.09.22.508985

**Authors:** Christoph Witzel

## Abstract

Negative colour afterimages incorporate the most fundamental tenets of human colour perception (adaptation, opponency, complementary colours, context-dependence, dependence on the beholder), but despite their fundamental importance, the underlying mechanisms are controversial. The key question is whether the afterimages are attributable to adaptation in the cone photoreceptors, of colour-opponent neurons in the subcortical pathway, or require the assumption of yet unknown cortical mechanisms. The most common assumption in textbooks and contemporary research is that negative afterimages are cone-opponent. It has not previously been recognised that the role of those mechanisms can be distinguished because they make fundamentally different predictions about the hue and saturation of afterimages. To test these predictions, we developed experimental paradigms to measure the exact colours perceived in afterimages. The results reveal that afterimages do not align with cone-opponency but closely follow a well-founded model of cone adaptation (cone contrasts). Our findings establish that cone-adaptation is the sole origin of negative colour afterimages. The quantitative cone-contrast model provides a comprehensive, straight-forward explanation of the adaptive mechanisms underlying colour afterimages that resolves apparent contradictions and debunks wide-spread misconceptions. This model has far-reaching implications for longstanding mysteries about visual perception.

## Summary / Introduction

Negative colour afterimages are illusory colour experiences that occur after fixating and adapting to a coloured area for a sustained time^1^. Such afterimages incorporate the most fundamental tenets of human colour perception: Chromatic adaptation, opponency, complementarity, and the dependence on context and beholder^2-8^. The adaptation underlying the induction of afterimages has been attributed to different mechanisms at all stages of the visual hierarchy. The body of research on afterimages is utterly contradictory, either confirming or refuting that afterimages are the result of adaptation of the cone photoreceptors (bleaching)^9-12^, of the cone-opponent channels^2,5,13-17^, of Hering-opponent^5,8,18^, or other cortical mechanisms^15,19-28^. The most wide-spread assumption in textbooks^3-5,7,8^ and contemporary research^13,29^ is the idea that cone-bleaching produces cone-opponent afterimages. Empirical inconsistencies with that idea have been attributed to additional, yet unknown cortical mechanisms^5,19-28^. Here, we show unequivocal evidence that afterimages originate from photoreceptor adaptation, resolving all misconceptions and contradictions. Using new paradigms, we measured the precise hue and saturation of afterimages for a wide range of inducers. We quantitatively predict the colours perceived in afterimages through a well-founded model of cone bleaching (cone contrasts)^30,31^. Results demonstrate that the colour of afterimages tightly matches the non-opponent predictions of the model.

## Results

Research and textbooks overlook the fact that the adaptation of different sensory mechanisms makes fundamentally different predictions about the hue and saturation of afterimages^2-8,16,29^. Cone-bleaching is incompatible with colour opponent afterimages, and neither cone-bleaching nor adaptation of the cone-opponent channels are compatible with Hering’s colour-opponency (Figure 1). These predictions can be tested by experimental paradigms that measure the exact colours perceived in afterimages. We developed two such paradigms, a fixed-location and a chaser-like simultaneous matching task (Figure 2.a & e).

**Figure 1.**
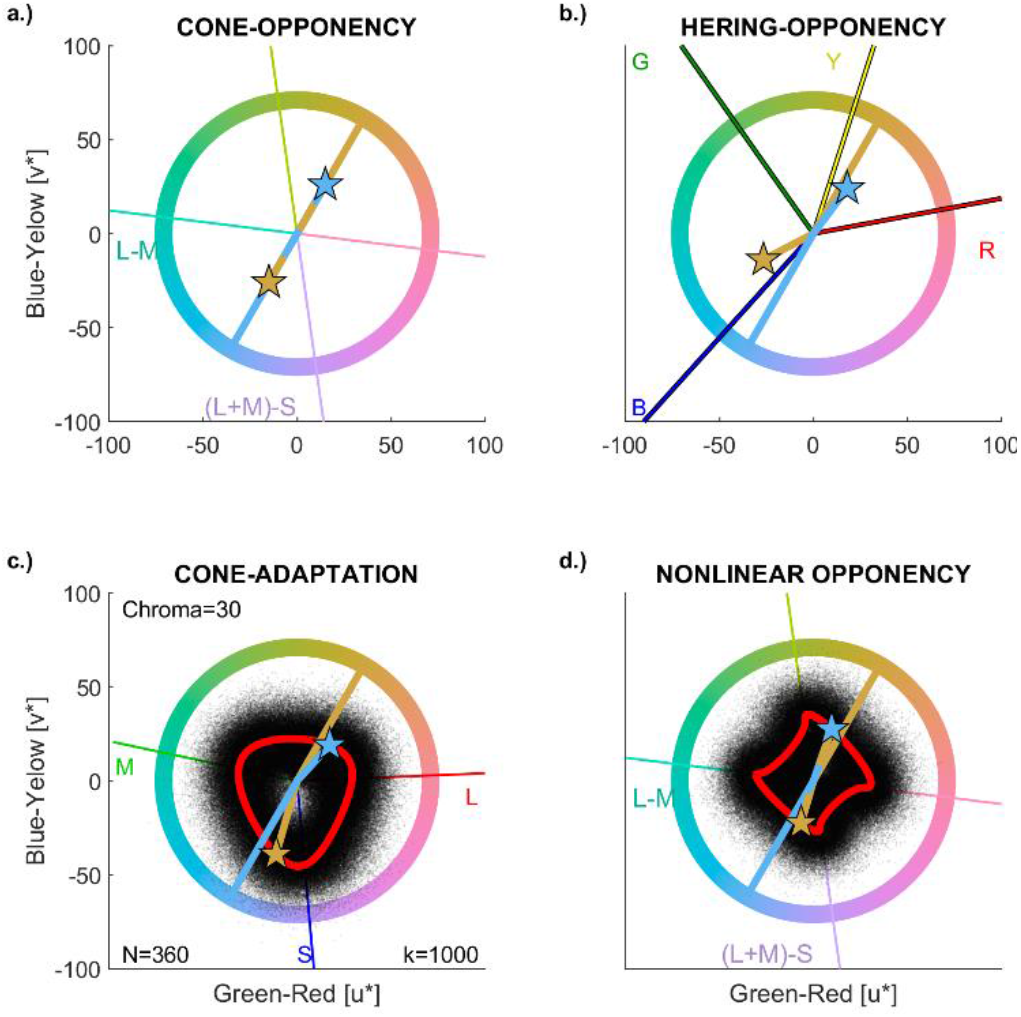
Afterimage Models. Panel a illustrates linear adaptation of the cone-opponent channels, where the afterimage is shifted by a proportional chroma to the hue opposite to the inducer hue. Panel b illustrates Hering opponency. The coloured lines in the background correspond to measured prototypes of red, yellow, green, and blue. Panel c shows the results of the cone-contrast model without (red line) and with noise (black dots). Panel d exemplifies nonlinear adaptation along the cone-opponent channels. Note the difference of predictions in panels c-d from a circle that would result when afterimages were cone-opponent. Two opponent examples of inducers are shown. The crossing point between line and circle indicates the inducer, stars show the colour of the corresponding afterimage predicted by the respective model. Line and stars are shown in the inducer colour. For first-person inspection, the yellowish inducer (60 deg) corresponds to the inducer in Animation1 in the supplementary material.

**Figure 2.**
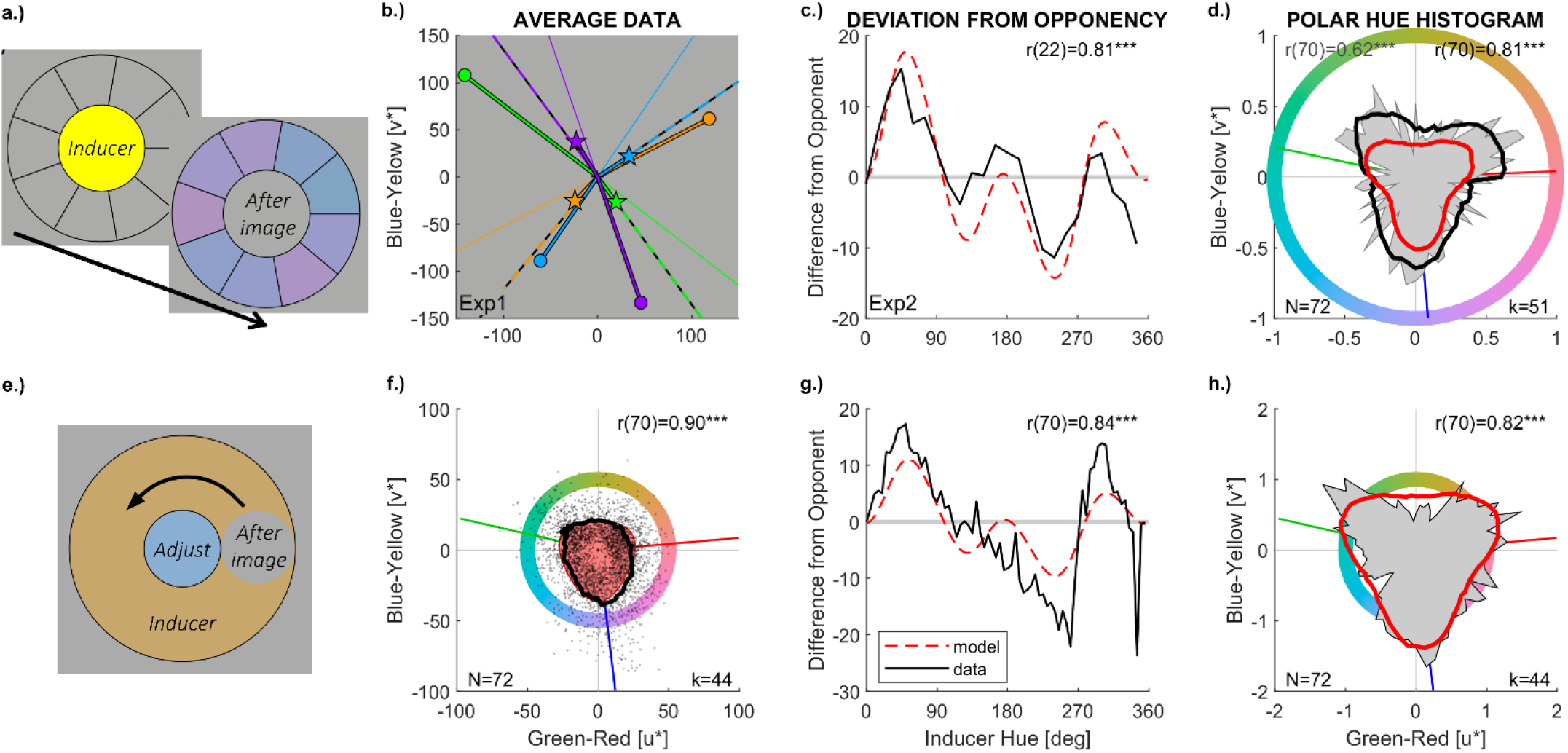
Tasks and Results. The first row (a-d) illustrates stimulus displays and task (a), and results (c-d) for the first experimental paradigm. Observers adapt to the inducer display for 30s (yellow in a). Then the inducer display is replaced by a circle of background grey together with nine comparison colours (purple in a) in a circular arrangement around the grey circle. The grey circle in the centre is perceived as the induced afterimage, and observers select the comparison colour that best matches the colour they see in the centre. Panel b gives examples of afterimage matches from Experiment 1 with maximally saturated colours. Discs correspond to inducers, stars to average matches of perceived afterimages. The thin line in the background indicates the cone-opponent direction, the dashed line indicates the prediction of the cone-contrast model (See Figure S1 for results with the other four colours). Panel c-d illustrate results of Experiment 2. The curves in the third column show the deviation of the cone-adaptation model (dashed red curve) and of the measured afterimages (black curve) from the colours opponent to the inducers. The correlation reflects the high similarity in profile of the two curves. Correlations between simulated and measured afterimage hues are shown at the top of the diagrams. The second row (e-h) corresponds to the chaser-like paradigm. In panel e, the chromatic ring is the inducer. Participants fixate the centre, and the moving grey circle on the ring produces the afterimage. Observers adjusted hue and chroma of the centre circle to match the moving one. Average afterimage matches (black curve in f) closely correspond with model predictions (red area in the background), yielding a correlation (top right) between measured and simulated afterimage intensity (i.e., chroma). The polar histograms in the last column (d,h) counts hue responses (azimuth in panel f) and displays the resulting frequencies as a function of azimuth in a polar plot. The histogram of the measurements is shown by the grey area in the background, the smoothened version of that histogram by the black line, and the histogram of the simulated afterimages by the red line. Polar hue histograms feature three clusters that closely correspond with model predictions (red line), hence confirming the results from panels c, f, and g. The attached Animation1 illustrates the pronounced difference between the afterimage and the cone-opponent hue of the yellowish inducer in panel e: The afterimage (which appears after staring at the centre for several seconds) is much more purple than the cone-opponent colour (which is displayed in the animated illustration when the moving circle is in the bottom-right quarter of the inducing ring). Animation2 illustrates the clustering of afterimages from different inducers.

### Fixed-location Task

The first task combined a typical afterimage display with simultaneous matching (Figure 2.a). Observers simultaneously matched the hue of comparison stimuli with the hue of an afterimage that appears at the location of an inducing colour after that inducing colour disappeared. Chroma of comparison stimuli was kept constant (see Method). In a first experiment, measurements with highly saturated colours showed that afterimage hues were neither cone-opponent to the hues of the inducers (Figure 2.b), nor opponent according to Hering’s green-red, and blue-yellow axes (Figure 1.d), nor according to other models of colour appearance (Table S1-2). The deviations of the measured afterimage hues from cone-opponent hues were almost perfectly correlated with predictions of the cone-contrast model (r(6) = .98, p < .001). These strong effects on hue selection were specific to afterimages, and did not occur in a control task, in which observers matched real colours in an otherwise same test display (Figure S1).

We ran a second experiment to test the above correlation with a larger stimulus sample. Twenty-four inducers were sampled at equal intervals from an isoluminant, equally saturated circle in CIELUV space (the coloured circle in Figure 1). If afterimages were attributable to proportional adaption of cone-opponent mechanisms the afterimage hues would be equally spaced around a circle (see Method). If there was non-linear adaptation to cone-opponency or any other kind of colour opponency (e.g., Hering colours), afterimage hues should align with the four directions of the respective two opponent dimensions (Figure 1.d). Neither prediction was supported. Instead, observers’ matches showed that the hues of afterimages were systematically shifted away from the cone-opponent hue directions (Figure 2.c) towards three directions that roughly coincided with the directions of maximal cone excitations (Figure 2.d). To make precise predictions, we modelled cone adaptation through cone-contrasts (red line in Figure 1.c). The deviations of the measured afterimage hues from cone-opponent hues were strongly correlated with predictions of the cone-contrast model (r(22) = .81, p < .001; Figure 1-c). To simulate a hue histogram, we added response noise (black dots in Figure 1.c) to obtain a continuous probability distribution (black curve in Figure 2.d). This cone-adaptation model predicted the frequencies of hue selections with a correlation of over r(70) = .62, p < .001, across the 72 response hues. The distribution of responses for 24 inducers across 72 response options produced artefactual variation (zigzags in the grey area of Figure 1.c) that undermines a correlation with the probability distribution from the model. Smoothening the distribution (red curve in Figure 2.d) counteracts these artefacts and yielded a correlation with the model of r(70) = .80, p < .001. There was no such correlation with distances of inducer hues from cone-opponent (DKL) and Hering-opponent axes (all p < .37; Table S3).

### Chaser-Like Task

The model also makes predictions about the strength, i.e., the chroma, of the afterimages (Figure 1.b). We used a chaser-like task (Figure 2.e) to maintain the afterimage over time, allowing for adjustments of both, hue and chroma, to match the induced colours without decay of afterimage strength over time. The average adjustments across 72 inducers (black curve in Figure 2.f) closely followed the predictions by the cone-adaptation model (red area in Figure 2.f), resulting in a correlation of r(70) = .90, p < .001, between predicted and measured chroma. As in the other task, deviations from cone-opponent hues were correlated between simulated and measured afterimages (r(70) = .84, p < .001, Figure 2.g), and the hue histogram for measured and modelled afterimages were strongly correlated (r(70) = .82, p < .001; cf. Figure 2.h). These results could also be produced at the individual level: For each of the 10 observers, there was a correlation between predicted and measured chroma (r(70) = .42 to .84, all p < .001) and between measured and simulated hue histograms (r(70) = .27 to .61, all p < .05); and for 9 out of 10 there was a positive correlation between measured and simulated deviations from cone-opponent colours (r(70) = .35 to .90, all p < .01, except: r(70) = 0.22, p = .07). The strength of afterimage induction through yellowish inducers and the shift of the corresponding afterimages away from cone-opponent blue towards the purplish S-cone direction are so pronounced that they are obvious to first-person experience (see attached Animation1: the afterimages would be blue according to cone-opponency like the colour appearing in the centre but is actually purple).

The cone-contrast model also predicts nonlinear changes in hue when increasing inducer chroma (Figure 3.a). Afterimages have been measured across 3 different chroma levels for 24 hues with observer cw. Differences from opponent colours were strongly correlated between measurements and simulations based on the cone-contrast model (r(70) = 0.78, p < .001), reproducing the above observations for measurements across chroma (Figure 2.c,g). To specifically test the change of hue across chroma, we calculated, for each inducer hue, the difference between the afterimage hue measured at each chroma level from the average afterimage hue across chroma levels (Figure S5 for illustration, Figure 3.b for results). These chroma-specific hue differences were strongly correlated when calculated between cone-contrast predictions and measurements (r(70) = 0.65, p < .001), showing that perceived hue changes with chroma in the way predicted by the model.

**Figure 3.**
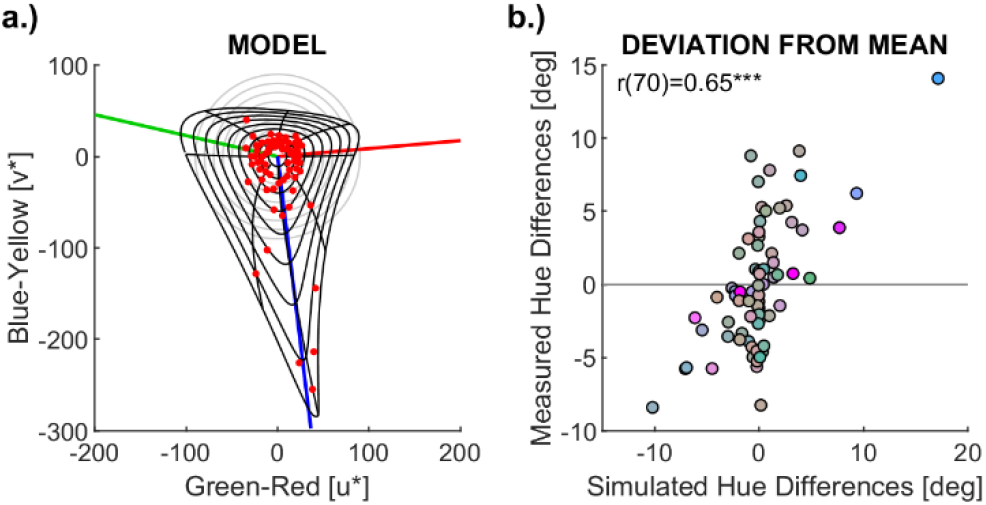
Change of Afterimage Hue with Chroma. (a) Afterimages modelled through cone contrasts. The grey lines in the background illustrate inducer colours at different levels of chroma (radius); the black lines illustrate how the corresponding afterimages vary with chroma. Red dots indicate simulated afterimages for measurements across different levels of chroma. (b) Scatterplot that illustrates the correlation between simulated and measured changes of afterimage hue across chroma. The axes indicate the afterimage hue difference from the average hue across chroma levels separately for each inducer hue. Data for observer cw.

## Discussion

All results of the two different afterimage-inducing paradigms confirm that afterimages do not align with cone-opponent mechanisms but follow instead a well-founded model of cone adaptation. The cone-contrast model explains large parts of the variance in hue (>60% of variance) and chroma (>80%) of afterimages (Figure 2). Despite variation across observers, almost all individual data provide unequivocal evidence for the signature of cone-adaptation when accounting for individual differences in induction strength in the model (Figures S2-4). Hue changes of afterimages across chroma also followed the cone-contrast model (Figure 3). Considering measurement noise, our results suggest that afterimages are fully determined by cone-bleaching at the first stage of colour processing.

Such afterimage formation can be explained through a time lag between cone bleaching and regeneration^9,32^. The cone-adaptation model used here implements adaptation according to Weber’s Law, which implies that cone sensitivities decrease by decrements that are proportional to the magnitude of cone-adaptation. This model reflects the proportional decrease of photon catches due to cone bleaching: a given light has a certain probability with which it isomerises molecules of retinal bound to opsin. When retinal molecules already underwent isomerisation due to ongoing stimulation by an inducer, that probability applies to the remaining proportion of retinal molecules, implying that the amount of isomerising retinal molecules is proportional to the available, non-isomerised molecules^30,31^. As a result, the adapted cone response is proportional to the inverse of the molecules bleached (isomerised) by the inducer. This multiplicative inverse relationship with inducer cone excitation produces non-linear effects of cone adaption, resulting in the three peaks of afterimage strength, reflected by afterimage chroma and the three hue clusters.

At high saturation, each one of the three afterimage peaks is determined by the single most strongly adapted cone (Figure S6). At low saturation, they result from by the least adapted cone, i.e., the other two most strongly adapted cones: If L- and M-cones are most adapted by inducers, S cones dominate the afterimage; if L- and S-cones are adapted, M-cones dominate the afterimages; and if S- and M-cones are most adapted, L cones dominate the afterimages. L- and M-cone sensitivities overlap. So, maximal adaptation of one always implies some adaptation of the other, and the three local maxima (peaks) of the cone model do not coincide with the directions of isolated cone excitations (red, green, and blue lines in Figures 1.c). The overlap between L- and M-cones is stronger for broadband spectra of desaturated colours producing a shift of chroma away from isolated cone excitations with decreasing chroma (bent black lines in Figure 3.a). Since S-cone sensitivity barely overlaps with M- and L-cone sensitivity, the effect of S-cones on L-M-adapted afterimages is much stronger than the effects of the correlated L-or M-cones, resulting in a pronounced asymmetry along the cone-opponent S-(L+M) axis (close to the blue-yellow axis v*, cf. Figure 1.b and 3.a). The clustering of afterimage hues resulting from the nonlinear relationship between inducers and afterimages, implies that visibly different inducers may produce very similar afterimages, as illustrated by the attached Animation2.

Cone-bleaching had been a classical explanation of afterimages^9,12^, but was refuted by observations that afterimages were not equivalent for inducers that had, on average, equal intensity^10,11^. This contradictory evidence assumed that cone-bleaching and regeneration would cancel each other over time, assuming a linear relationship of cone-bleaching and pigment regeneration over time. However, this assumption does not hold. The non-linear, multiplicative inverse relationship between cone-excitation and resulting cone-adaptation is also found in the time course of cone bleaching and regeneration^33^, and in the time course of chromatic adaptation^34^. In addition, the flicker may be triggering adaptive mechanisms beyond the photopigment kinetics that are not specific to afterimages. So, the observation that different temporal sequences of inducer stimulation produce dissimilar afterimages^10,11^ is fully compatible with our findings.

Most state-of-the-art research assumes that afterimages correspond to the colour predicted by cone-opponent mechanisms at the second stage of colour processing^2,5,13,14,18^, but again others found contradictory results^15-17^. Evidence for this idea has been provided by tasks showing the successful nulling of afterimages in cone-opponent space^13^. However, successful nulling works independently of whether the appearance of afterimages follows a straight line due to second-stage, cone-opponent adaptation, or a curve due to nonlinear adaptation, such as cone bleaching. Instead, our findings explain why afterimages are not cone-opponent, not reciprocal, and produce three clusters of hue (cf. Figure 1.c).^16^

Unlike what current textbooks suggest^3-5,7,8^, colour afterimages are neither determined by cone-nor by Hering-like colour opponency. A cursory look at afterimages may be misleading because afterimages resemble cone-opponency at a very coarse level^29^. This is not due to adaptation of the cone-opponent mechanisms at the second stage of colour processing, but to the cone-adapted colour signal from the first stage being propagated to the second and subsequent stages. This propagation of the afterimage signal also explains the physiological responses of retinal ganglion cells as a function of afterimage decay^13^. The present experimental paradigms allowed a closer look at the precise colours of negative afterimages and reveal that afterimages are not aligned with cone opponency but are almost fully determined by first-stage adaptation. Little variance of perceived afterimages is left that would support additional adaptation at later, subsequent levels of processing. Summoning magic (i.e., unknown and omnipotent) cortical mechanisms to explain deviations from cone-opponency is unnecessary.^15,28^

Still others, claimed that afterimages occur through adaptation of cortical mechanisms that involve shape and object recognition^19-22,35^, binocular integration^23,24^ and/or colour constancy^25^, contradicting classical evidence for a retinal origin^26,27^. Top-down effects on colour perception are well known^36,37^. Such effects may be particularly strong for illusory percepts like afterimages where a physical stimulus is absent^22^. However, evidence for mid-level (e.g., contours and shape^19,22^) or high-level (e.g., knowledge^21,35^) effects on afterimage do not necessarily reflect the origin of afterimages. Afterimages occur in the absence of top-down cues or knowledge, as in the present experiments. Hence, top-down interference effects may modulate the subjective appearance of afterimages; but they are not the origin of afterimage formation.

From the present results, it is clear that negative colour afterimages are determined by cone adaptation and can be precisely predicted by a straight-forward cone-contrast model of the first stage of colour processing. These findings debunk the most widespread misconception about the cone-opponency of complementary afterimages, including approaches based on RGB, HSV, and xyY, which all mathematically imply cone-opponency^**3-5**,**7**,**8**^. Yet, our findings do not only decide over general ideas in a decades-old debate on the origin of afterimages. They establish a quantitative model that comprehensively explains afterimage formation in detail. The mechanisms of adaptation underlying afterimage formation play a fundamental role in visual perception, determining the ability to discriminate colours and the way colours subjectively appear to human observers^7^. The quantitative model of afterimages provides new ways of investigating a whole range of yet ununderstood visual phenomena (cf. Section D of the Supplementary Material). For these reasons, clarifying the mechanisms underlying afterimages is fundamental for understanding colour perception and for the measurement, specification, and control of colours (colorimetry) in scientific and industrial applications^**3-5**,**7**,**8**^. It’s time to update textbooks.

## Method

### Task 1: Fixed-Location Afterimages

#### Participants

In Experiment 1, 32 observers participated (20 women, age: 25.2±3.94y). In Experiment 2, 52 observers took part (36 women, age: 25.1±4.3years). Participants were compensated by course credits or €8 per hour. Colour vision deficiencies were excluded with the HRR plates^38^. The experiments were approved by the ethics committee at the University of Gießen (LEK 2017-0030), and informed consent was obtained from all observers.

#### Apparatus

Stimuli were presented on an Eizo Color Edge Monitor (36,5 × 27cm) with an AMD Radeon Firepro graphics card with a colour resolution was 10 bit per channel. The CIE1931 xyY chromaticity coordinates and luminance of the monitor primaries were *R* = (0.6851, 0.3110, 27.1), *G* = (0.2133, 0.7267, 69.4), and *B* = (0.1521, 0.0450, 4.7). Gamma was 2.2 for all channels and has been corrected.

#### Stimuli

Colours were represented in CIELUV space. The white-point was [0.3304, 0.3526, 101.1], background lightness was at L*=70. At isoluminance, opponent hues in CIELUV are the same as cone-opponent hues in DKL-space (Figure 1.a) and in (gamma-corrected) HSV-space. CIELUV-space was preferred to DKL-space because it better controls for perceived chroma^39^. The eight inducer colours in Experiment 1 were chosen to correspond to the typical lightness and hue of red, orange, yellow, green, turquoise, blue, purple, and magenta at the maximum chroma possible within monitor gamut. The nine hues of the comparison colours were obtained by adding four hues in 10 deg steps to either direction (low or high azimuth) of the opponent hue. Lightness and chroma of the comparisons were determined through piloting. In Experiment 2, twenty-four inducers were sampled along a hue circle in CIELUV at chroma 71 and equal steps of 15 deg starting at 0 deg (cf. hue circles in Figure 1). Chroma was chosen to be the highest chroma achievable within monitor gamut for all hue directions. Lightness of inducers, comparison colours, and the achromatic disc in the centre of the test display were the same as the background (L* = 70). Chroma of all comparison colours was kept constant at 30. This level of chroma was determined through piloting and accounts for the lower saturation of the afterimages as compared to the inducers. Table S1 provides detailed colour specifications.

#### Procedure

In one trial, the inducer display was shown for 20 seconds (Experiment 1) and 30 seconds (Experiment 2). During this time, a round fixation dot was blinking at a rate of 1Hz at the center of the screen. Observers were asked to stare at this dot. To make sure they would, they also completed a cover task. In some trials, the round dot would change for one blink into a square dot. Observers had to indicate at the very end of a trial whether such a squared dot occurred in the trial or not.

While seeing the afterimage, observers determined which of the nine comparisons was the best match. Afterimages tended to melt into segment with the colour that looked closest to the afterimage, making the task intuitive. Participants used the mouse to indicate the segment with the matching comparison (Figure 2.a). If they could not see any of the comparison colours in the centre, they could also click on the centre disk to indicate this (resulting in a missing value). After this response, participants responded to the cover task, and if wrong, had to redo the whole trial at the end. A 10-seconds intertrial display followed by a self-parsed break was used to cancel remaining afterimages and to prevent afterimage interference across trials (see Section A of Supplementary Material).

### Task 2: Chaser-Like Afterimages

#### Participants

Ten voluntary participants (6 women), including the author (CW) took part (cf. Table S4, Figures S2-4). Measurements across chroma were done by the author (CW). The experiment was approved by the Ethics Committee at the University of Southampton, ERGO 65442.

#### Apparatus & Stimuli

Three different **e**xperimental set-ups were used and calibrated as explained for Task 1 (see Table S4 for details). All measurements were conducted in ambient darkness. 72 inducer colours were sampled in 5-deg steps along an isoluminant hue circle at L*=70 in CIELUV-space. Inducer chroma was set to be maximal within the respective monitor gamut, resulting in a chroma of 38 (one participant), 42 (two participants), or 50 (everyone else). Figure 1.e and the attached Animation1 illustrate the stimulus display. For measurements across chroma (Figure 3), chroma levels at 20, 42, and maximum within gamut were measured for hues of 15 deg differences, starting at 0 degree.

#### Procedure

In each trial, participants were asked to fixate the centre of the display until the moving circle reaches maximum colourfulness. Then, they adjusted the hue (left/right) and saturation (up/down) of the centre circle using the cursor keys. During a coarse adjustment, they could continuously change colours; but before confirming the adjusted colour, participants needed to do a fine adjustment (by pressing space). In the fine adjustment, colour change by single steps (in radius or in 1 deg azimuth) with each separate key stroke.

Since the adjustment takes time, observers would see afterimages from the adjusted colour attenuating the chroma of the adjusted colour. Two measures were taken to avoid this: During coarse adjustment, black and white circles (not areas) were extending from the centre of the adjusted disks to its rim. These circles would disappear during fine adjustments to avoid interference with the finalised match. In addition, pressing the control key would turn the centre disk temporarily into grey until key release. Observers were asked to wipe out unwanted afterimages from the adjusted centre disk using that key before confirming the adjustment. So, they press control, wait and move their eyes until the centre disks looks achromatic. They then stare at the grey circle until the moving circle reaches maximum chroma. Only then, they released the key and compared the colour they had previously adjusted with the colour of the moving circle. Typically, the adjusted colour was too saturated due to the overlay of the self-afterimage during the adjustment. So, participants needed to lower chroma. They reiterated this procedure until the chroma did not need adjustment after releasing the control key. Only then, participants confirmed adjustments by pressing enter. As in Task 1, the intertrial period was used to prevent afterimage carry-over across trials.

Inducer colours were split into nine series of eight colours. Within a series, the eight colours differed by 45 degrees azimuth. Different series were defined by different starting points (0, 5, 10…40 degree) so that the series covered all 72 inducers. One block of measurements featured one trial for each of eight inducer colours in a series. Measurements of each block were repeated up to five times (cf. Figure S2-4). Blocks of measurements were spread over several days. Prior to measurements, participants were trained with practice blocks in the presence and with the feedback of the experimenter (CW) to make sure they understood handling and its purpose.

### Cone-Adaptation Model

Adaptation and cone bleaching at the first stage of colour processing were modelled by cone contrasts^40-42^. Cone contrasts (CC) are Weber fractions, calculated as the difference between cone excitations of the stimulus (CE) and cone excitations of the adapting colour (CE0) relative to the cone excitations of the adapting colour: CC = (CE-CE0)/CE0. These cone contrasts model cone bleaching^30,31,43-46^. The reduction of sensitivity at high stimulus intensities can be explained by the reduction of available photopigment due to bleaching. Since bleaching is proportional to intensity of the adapting stimulus, one needs proportionally more stimulation (likelihood of photon catches) for detecting a non-adapted stimulus. In psychophysical experiments, adaptation is typically controlled by the colour of the background. In the case of afterimages, the roles are swapped because we model perception of the achromatic background after local adaptation to the inducer. Hence, CE0 is the cone excitation of the inducer, and CC the one of the background. Cone contrast changes with increasing cone adaptation CE0 according to a multiplicative inverse function (because CE0 increases in the denominator). Cone contrast is calculated independently for the short-(S), medium-(M), and long-wavelength (L) sensitive cones, resulting in S-cone, M-cone, and L-cone contrasts. Cone adaptation produces shifts towards the peak cone sensitivities, as illustrated by the three peaks in a hue histogram (Figure 1.c, Figure 2.d,h).

The observation that afterimages are not as saturated as inducers (Figure 2.f) shows that adaptation to inducers is not complete. As an estimate of the strength of adaptation, we set the adapting chroma for modelling afterimages to half the chroma (35.5 deg) of the inducers in the first task, and to the grand average chroma (27.0 deg) of adjustments for the second task (which is slightly more than half the average inducer chroma, 23.3 deg). Even if we assumed complete adaptation to the chroma of inducers, results would be largely the same. However, not surprisingly, correlation coefficients would be slightly lower because non-linearities in the model are then higher than in the measurements. There is no free parameter in this model.

## Supporting information

AnimatedIllustration

## Acknowledgements

Thanks to Bevil Conway, Daniel Osorio, Karl Gegenfurtner, Kevin O’Regan, Matteo Toscani, Mike Webster, Shin’ya Nishida, and Wendy Adams for helpful comments, and Alexander Nowak for help with data collection. Research was partially funded by the Deutsche Forschungsgemeinschaft (DFG, German Research Foundation) – project number 222641018 – SFB/TRR 135 TP C2, and research funds from the School of Psychology at the University of Southampton.

