## Supplementary figures and images for "The Mechanisms Underlying Colour Afterimages"

### AnimatedIllustration

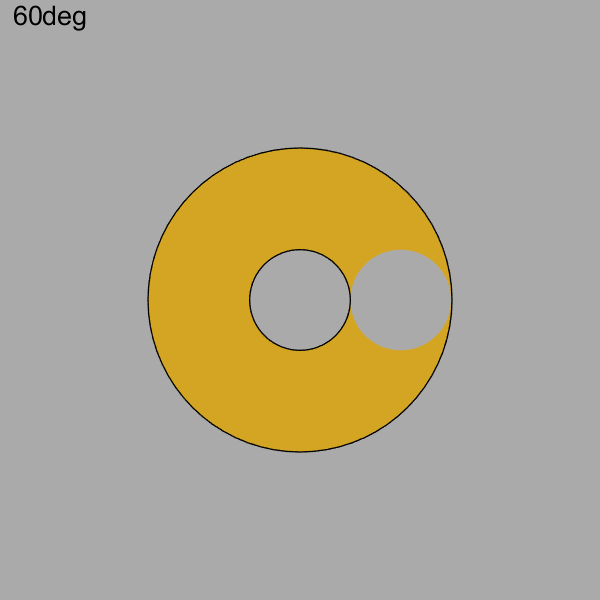
